# Rate of force development is correlated with corticospinal excitability during explosive voluntary contractions

**DOI:** 10.1101/2024.08.27.607589

**Authors:** Federico Castelli, Omar S. Mian, Adam Bruton, Ashika C. Chembila Valappil, Ricci Hannah, Neale A. Tillin

## Abstract

**Objective:** To investigate the relationship between rate of torque development (RTD) and corticospinal excitability (denoted by motor-evoked potential; MEP) during explosive voluntary contractions. Also, to assess differences in MEP and silent period duration between different phases of explosive contraction and at maximum voluntary contraction (MVC) plateau.

**Methods:** In 14 adults, quadriceps muscle MEP and silent period duration were measured at ∼45 (early), ∼115 (middle), and ∼190 ms (late) from EMG onset during knee-extensor isometric explosive contractions, and at MVC plateau with superimposed transcranial magnetic stimulation (TMS). RTD was measured immediately prior to early-phase MEP during the same contractions, and these two variables were correlated across separate contractions, within participants, via repeated measures correlation (RmCorr). RTD was also measured in explosive contractions without TMS over early-, middle-, and late-phases and correlated to MEP averaged across the corresponding and preceding phases, via Pearson’s r correlations, assessing relationships across participants. MEP and silent period duration were compared (ANOVA) between the different phases and at MVC plateau.

**Results:** MEP and RTD were correlated across separate contractions within participants (RmCorr r=0.43), and in the middle phase across participants (Pearson’s r=0.56). MEP and RTD were not correlated in other phases (r≥0.09). Silent period duration increased throughout the different phases of contraction and up to MVC plateau (ANOVA, p=0.001) but MEP remained constant (ANOVA, p=0.42).

**Conclusion:** The correlations between MEP and RTD suggests corticospinal excitability is an important determinant of RTD. During rapid torque development and up to MVC plateau, corticospinal inhibition increases but excitability remains constant.

## Introduction

The ability of muscles to rapidly increase joint torque, measured as the rate of torque development (RTD), is important for movements where the time to develop torque is limited, such as sprinting (Tillin, Pain and Folland, 2013) or balance recovery (Sundstrup *et al*., 2010). RTD is often measured in early (0-50 ms), middle (0-100) and late (>100 ms) phases of explosive contraction performed from rest (Maffiuletti *et al*., 2016). Maximum voluntary contraction (MVC) torque is an important determinant of late-phase RTD (Folland, Buckthorpe and Hannah, 2014), while muscle activation, determined by the number of motor units recruited and their discharge rates, is an important limiting factor of early and middle phase RTD (Folland, Buckthorpe and Hannah, 2014; Del Vecchio *et al*., 2019). Despite the relevance of muscle activation to RTD, corticospinal behaviour during explosive contractions and its association with RTD is not well established. The corticospinal mechanisms affecting RTD could be explored using transcranial magnetic stimulation (TMS), superimposed during the different phases of explosive voluntary contractions. Specifically, superimposed TMS elicits motor evoked potentials (MEP) and silent period durations immediately after the MEP, in the EMG signal of the target muscles, which are thought to reflect corticospinal excitability (Rossini *et al*., 2015) and inhibitory mechanisms (Säisänen *et al*., 2008; Yacyshyn *et al*., 2016), respectively.

Compared to slow (low RTD) ramped contractions, explosive (high RTD) contractions are characterised by lower motor unit activation thresholds (Desmedt and Godaux, 1977) and high motor unit discharge rates (Van Cutsem, Duchateau and Hainaut, 1998; Duchateau and Baudry, 2014; Del Vecchio *et al*., 2019). This high muscle activation seems necessary to maximise RTD, a theory further supported by observations of positive correlations between agonist EMG amplitude in the early phase (0-50 ms) of EMG activity – which commences prior to torque onset (Tillin et al., 2010) – and early to middle phase RTD (Folland, Buckthorpe and Hannah, 2014). To achieve high muscle activation (and thus high RTD), it is conceivable corticospinal excitability will need to be high, given that corticospinal excitability increases with contraction intensity (Todd, Taylor and Gandevia, 2003; Goodall, Romer and Ross, 2009; Weavil *et al*., 2015). We might, therefore, expect RTD and MEP amplitude to be positively correlated in the early- to middle phases of contraction, both within a person across separate contractions and across participants.

In addition to high corticospinal excitability, low corticospinal inhibition during the early phase of EMG activity may be necessary to optimise RTD. Cortical inhibition, measured by reduced short-interval intracortical inhibition, appears to decrease in preparation for an explosive contraction immediately prior to EMG onset (Baudry and Duchateau, 2021). Should low inhibitory conditions persist after the onset of EMG activity during an explosive contraction, and should this low inhibition be necessary to optimise RTD, we may expect the TMS-induced silent period duration to be short and negatively correlated with early and middle-phase RTD.

Immediately after the initial activation of motor units at the start of an explosive contraction, the motor unit discharge rate drops from up to ∼189 Hz to <50 Hz (Desmedt and Godaux, 1977; Van Cutsem, Duchateau and Hainaut, 1998; Del Vecchio *et al*., 2019), and remains this low at the plateau of an MVC (Pucci, Griffin and Cafarelli, 2006). Additionally, EMG amplitude increases rapidly in the early phase of explosive contraction, reaching nearly 120% of the EMG observed at MVC plateau after 75 ms, before declining to stabilise at 100% of MVC (Tillin, Pain and Folland, 2018). These results collectively suggest greater muscle activation at early compared to later phases of explosive contractions and MVC plateau. This dynamic modulation of muscle activation may be achieved through either: (i) corticospinal excitability starting high and decreasing and/or (ii) corticospinal inhibition starting low and increasing, throughout the different phases of explosive contraction and up to MVC plateau. To the best of our knowledge either (i) or (ii) have not been investigated in explosive condition, while a study of ballistic wrist movements, showed MEP amplitude increased prior to movement onset but declined after movement onset (Mackinnon and Rothwell, 2000). Further, the increase in MEP amplitude preceded an increase in EMG amplitude, characterised by greater MEP/EMG in the early compared to later phases of contraction, suggesting corticospinal excitability increases disproportionately with muscle activation in the early phase, but changes proportionately with muscle activation at later phases of rapid contraction (Mackinnon and Rothwell, 2000). It is unclear if this phenomenon observed for dynamic rapid wrist movements is present during the isometric explosive contractions used to assess RTD.

This study has two main aims. The first aim investigates the relationships between RTD in the different phases of explosive contraction and both MEP amplitude and silent period duration in the same phases relative to EMG onset. These relationships will be explored between contractions within people and across participants. We hypothesise that MEP amplitudes in the early and middle phases will be positively correlated with RTD, whilst silent period durations will be negatively correlated. The second aim investigates the possible differences in MEP amplitude and silent period durations between different phases of explosive contraction (early, middle, late) and MVC plateau. We hypothesise that MEP amplitudes (absolute and relative to background EMG) will be greater and silent period durations shorter in the early phase of explosive contraction than in the other contraction phases.

## Methods

### Participants

Fourteen participants, nine males (age 31 ± 5 years, height, 178.7 ± 6.6cm, and mass 79.2 ± 4.7kg) and five females (age 29 ± 6 years, height, 165.5 ± 5.5cm, and mass 59.3 ± 6.2kg) were recruited to take part in this study. All participants habitually performed 120-180 minutes of moderate to high-intensity activity per week and were deemed recreationally active (McKay *et al*., 2022). Participants were also free from injury and disease (screened via a questionnaire adapted from Balady *et al*., 1998) and free from contraindications to TMS (screened via a questionnaire adapted from Rossi *et al*., 2011). Due to changes in endogenous hormones throughout the menstrual cycle potentially affecting neuromuscular responses to TMS (Ansdell *et al*., 2019), female participants undertook experimental trials exclusively during their self-reported early to mid-follicular phase (first ten days from the first day of menstruation), when endogenous hormone concentration is low and relatively stable (de Jonge, Thompson and Han, 2019). The University of Roehampton ethics committee approved the study, and all participants provided written informed consent before participating.

### Overview

Participants visited the laboratory on three separate occasions and were asked to avoid strenuous exercise and alcohol consumption for 24 hours before each visit. Each session lasted approximately 120-150 minutes, with consecutive sessions separated by 3-7 days. The first visit was a familiarisation session, and the second and third were measurement sessions. The measurement sessions were performed at a consistent time of day and involved an identical protocol. Repeat measurements could then be averaged across the two sessions to improve the reliability of the variables of interest. Sessions involved knee extensor torque and EMG measurements during MVCs and explosive voluntary contractions. TMS was used to obtain superimposed MEP and femoral nerve stimulation to obtain compound muscle action potentials at rest.

### Torque measurements and surface electromyography (EMG)

Participants were tightly secured in a custom-built strength testing chair (Fig.6b in Maffiuletti *et al*., 2016) with a waist belt and shoulder straps. The hip and knee angles were set at 100° and 105°, respectively (full extension being 180°). All contractions were isometric knee extensions performed with the right leg. An ankle strap joined to a calibrated S-shaped load cell (FSB-1.5kN, Force Logic, Reading, UK) was secured 4 cm proximal to the medial malleolus. The force signal was amplified (x375) and then sampled at 2000 Hz (Micro3 1401 and Spike2 v.8; CED., Cambridge, UK). Offline, the force was filtered (fourth-order low-pass Butterworth, 250 Hz cut-off), corrected for limb weight, and multiplied by the external moment arm to calculate joint torque.

The skin was prepared by shaving, cleaning (70% ethanol), and lightly abrading the area where EMG electrodes were placed. A single, bipolar silver-silver-chloride gel-electrode configuration (2-cm diameter and 2-cm inter-electrode distance; Dual Electrode, Noraxon, Arizona, USA) was placed over the belly of each of the rectus femoris (RF), vastus lateralis (VL) and vastus medialis (VM), based on SENIAM guidelines (Hermens *et al*., 1999). The three wireless EMG signals were transmitted to a desktop receiver (TeleMYO D.T.S., Noraxon, Arizona, USA) and sampled, along with a single wired EMG signal (see below), at 2000 Hz via the same A/D convertor and software as the force signal. The EMG system has an inherent 312-ms delay, so whilst suitable for measurements involved in the study, EMG signals sampled by it could not be used to detect activation onset and trigger the TMS in real-time during the explosive contractions (explained below in Experimental procedures). Thus, a second wired bipolar EMG electrode (2 cm diameter and 2 cm inter-electrode distance; Dual Electrode, Biometrics Ltd, Gwent, UK) was placed on the belly of the VM to trigger the TMS during contraction. Electrode locations were marked with a permanent marker pen, and participants were asked to maintain these marks throughout the study by re-applying them if necessary. Offline, wireless EMG signals were filtered (fourth-order Butterworth, band-pass, 6-500 Hz) and time-corrected for the 312-ms delay inherent in the Noraxon system.

### Transcranial magnetic stimulation (TMS)

TMS with a 1-ms pulse width was delivered via a double cone coil (110-mm Magstim 200, Whitland, UK) over the scalp in an optimal position to elicit MEPs in the right quadriceps muscles. The following procedures were completed in every session. Participants wore a swim cap, and the vertex of the head – identified as 50% of the distance between (i) nasion and inion and (ii) right and left temporomandibular joint – was marked on the swim cap. A 5-by-5-cm grid with 1-cm spacing between grid lines was drawn on the swim cap, lateral (left hemisphere) and posterior from the vertex. The coil was moved posteriorly and laterally from the vertex in ∼0.5-cm steps, and in each position, the participant completed four submaximal voluntary contractions at 20% MVC torque (established in the warm-up; see below) with superimposed TMS on each contraction at a submaximal (range 50-60%) stimulator output. The position which produced the highest consistent MEP amplitudes (peak-to-peak) over the four superimposed contractions for all three muscles (RF, VL and VM) was deemed the optimal coil position. Once established, this was marked by drawing the edge of the coil over the swim cap, which was used throughout the remainder of the session. The active motor threshold (AMT) was then determined via a series of 20% MVC torque contractions superimposed with TMS stimulator output starting at 39%. The AMT was defined as the minimum TMS intensity required to elicit five visible MEPs amongst the background EMG activity of the VM and RF out of ten consecutive superimposed contractions. If the muscles had less than five visible MEPs, the machine intensity was increased in steps by 2% of machine output (and conversely decreased if there were more than five visible MEPs). We prioritised the VM and RF (not the VL) in AMT decisions as these muscles produced more consistent and visible MEPs in pilot testing. Where it was impossible to match AMT for VM and RF, we settled on being one visible MEP away from 5/10 (e.g., 6/10 VM and 4/10 RF). The same investigator held the coil by hand throughout the measurement sessions, continuously monitoring its position and orientation. For TMS delivered during maximal and explosive contractions (see experimental protocol), the intensity was set at 140% of stimulator output at AMT. This intensity was chosen as it occurs on the steepest section of the input-output curve for MEP amplitude (Groppa *et al*., 2012; Rossini *et al*., 2015; Rossi *et al*., 2020) so it would maximise our chance of observing changes in this variable.

### Femoral nerve electrical stimulation

Single, square-wave pulses (200 μs duration) were delivered (DS7AH, Digitimer, Hertfordshire, UK) over the femoral nerve in the inguinal triangle to evoke twitch contractions and obtain compound muscle action potentials (M-waves) at rest. The anode (5 x 8 cm carbon rubber; EMS Physio Ltd., Oxfordshire, UK) was placed over the head of the greater trochanter. The optimal location of the cathode (1 cm diameter tip; S1 Compex Motor PointPen, Digitimer, UK) was determined as that which evoked the greatest peak twitch torque for a given submaximal stimulation intensity (80-120 mA). The cathode was taped down and held in position by the same investigator as a series of twitches at incremental intensities were evoked until there was a plateau in the peak-to-peak M-wave amplitude (Mmax) of all three muscles (RF, VL, VM). The stimulator intensity was then increased to 150% of the intensity at Mmax, ensuring supramaximal intensity. Three supramaximal twitch contractions were then evoked, each separated by 15 s, and Mmax was averaged across the three contractions for each muscle. The procedures were repeated for all sessions.

### Experimental protocol

Participants first completed a warm-up involving a series of explosive, submaximal, and maximal voluntary contractions (the latter being used to establish MVC torque), followed by procedures for obtaining optimal TMS coil position, ATM, and Mmax. Participants then completed a series of MVCs and explosive voluntary contractions with and without superimposed TMS. The instruction for MVCs was to “push as hard as possible” for 3-5 s and explosive contractions to “push as fast and hard as possible”, emphasising fast for 1 s. The protocol was organised into three blocks of contractions. Each block (Fig.1) involved 24 explosive contractions (8 without and 16 with superimposed TMS) and 8 MVCs (3 without and 5 with superimposed TMS) distributed across four sets. Each set involved six explosive contractions (the final four with TMS stimulation) and two MVCs. In sets 1-3, one of the two MVCs had superimposed TMS (randomly ordered), and in set four, both MVCs had superimposed TMS. Participants rested 10 s between explosive contractions, 30 s between MVCs, 120 s between sets, and 300 s between blocks. Each block was identical except for the timing of TMS application during explosive contractions, with each block using a different TMS timing for these contractions (see next paragraph). The order of the blocks was randomised across participants but held constant across sessions for the same participant. Overall, the 3-block protocol yielded 24 explosive contractions and 9 MVCs without TMS, 48 explosive contractions (16 per stimulus contraction phase) and 15 MVCs with TMS.

**Fig. 1.**
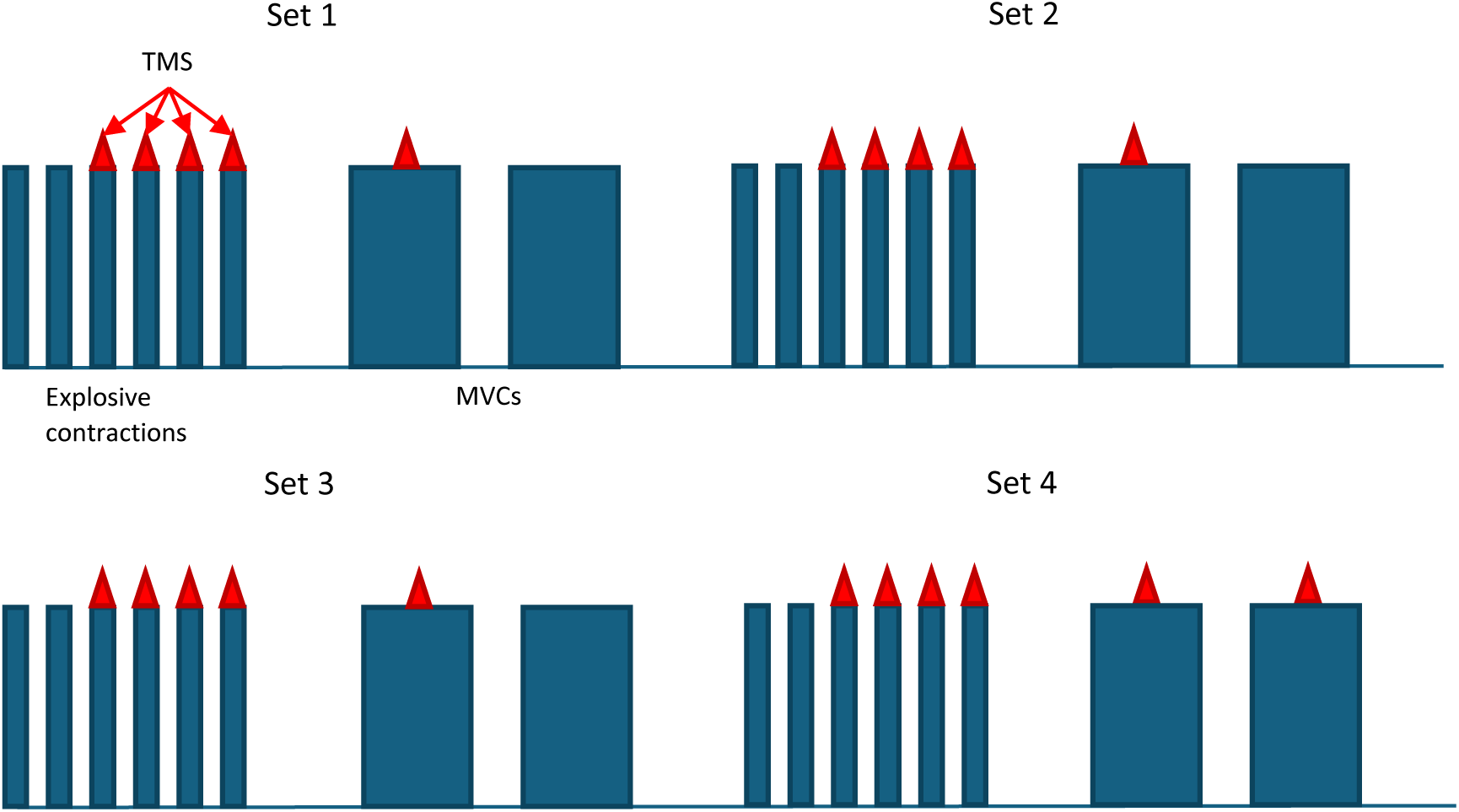
Schematic depicting a single experimental block, of which there were three per experimental session. Each block consisted of four sets of contractions, as shown. The thin blue bar represents explosive contractions, while the wide blue bar represents MVCs. The red triangle on the top of the blue bars represents the superimposed TMS. The blocks differed in timing of stimulation during explosive superimposed contractions, as described in the main text.

For superimposed MVCs, the TMS was triggered manually by the same experienced investigator during the torque plateau of the MVC. For superimposed explosive contractions, the TMS was triggered automatically when the VM EMG signal of the wired system exceeded a set threshold. The threshold was set in the Spike2 software as the lowest amplitude for that session above the highest peaks and troughs of the baseline noise. TMS was triggered at 3, 73, or 148 ms from EMG threshold crossing for early, middle, and late-contraction phases, respectively. When expressed relative to manually detected EMG onset (determined via the methods of Tillin et al., 2010), TMS triggers occurred at approximately 8 (early), 78 (middle), and 153 (late) ms. Thus, our threshold was typically crossed within 5 ms of manually detected EMG onset. This TMS triggering method was based on the methods of Giboin *et al*., 2018. The centre of the resulting MEPs occurred at approximately 45 (early), 115 (middle), or 190 ms (late) from manually detected EMG onset.

### Data screening and analysis

Out of 15 superimposed MVCs, only those where the TMS was delivered within 90% of MVC torque were used for further analysis. Out of the 16 superimposed explosive contractions per contraction phase, only those that met the following criteria were used for further analysis: (i) average baseline force did not change by >2 Nm during the 200 ms preceding manually detected force onset (detected as in Tillin *et al*., 2010); and (ii) there was a genuine attempt at an explosive contraction. A genuine attempt at an explosive contraction was defined as the instantaneous slope of torque-time, just prior to MEP onset, being within 3 standard deviations (SD) of the mean instantaneous slope at the same time point for contractions without TMS. The points for measuring instantaneous slope were 30, 105, and 180 ms after manually detecting torque onset for early-, middle- and late-phase contraction phases, respectively. Based on the above criteria, the first 10 useable contractions with TMS were used in each contraction phase for further analysis. This number was based on the following rationale: (i) our previous study (Castelli *et al*., 2024) showed averaging data across 10-20 contractions provided optimal reliability of the main dependent variables; (ii) the maximum number of usable contractions we could obtain in some participants in the early and middle contraction phases was 10.

For useable contractions, MEP amplitude was defined as the peak-to-trough of the superimposed response and is reported in absolute terms (V), normalised to Mmax (muscle-specific) and relative to muscle background EMG (MEP/EMG). Background EMG was RMS amplitude of the background EMG in the 20 ms immediately following TMS stimulation and therefore just prior to MEP appearance, within the same contraction and muscle as the MEP. After the MEP offset, the silent period duration was measured from the stimulation point to the EMG activity resumption. The second derivative of EMG amplitude over time was established (1-ms time constant) and then the signal was rectified to determine the resumption of EMG activity. EMG activity resumption was defined when the amplitude of the second derivative increased above 5 SD of the mean baseline (calculated in a 500 ms period prior to the contraction) for 70% of the next 10 ms (Damron et al., 2008; Fig.3). The semi-automated process was validated through manual inspection of the signal, as recommended by Damron et al., (2008). For correlations, TMS variables were averaged across the three quadriceps muscles as is normal when relating muscle activity of the quadriceps to net knee extension RTD (Tillin *et al*., 2010; Tillin, Pain and Folland, 2012, 2013, 2018; Folland, Buckthorpe and Hannah, 2014; Behrens *et al*., 2015; Morales-Artacho *et al*., 2018; Cossich and Maffiuletti, 2020). Torque at discrete time points from torque onset was used as a proxy for RTD (Maffiuletti *et al*., 2016), where torque onset was defined manually with a method considered the golden standard (Tillin *et al*., 2010).

**Fig. 2.**
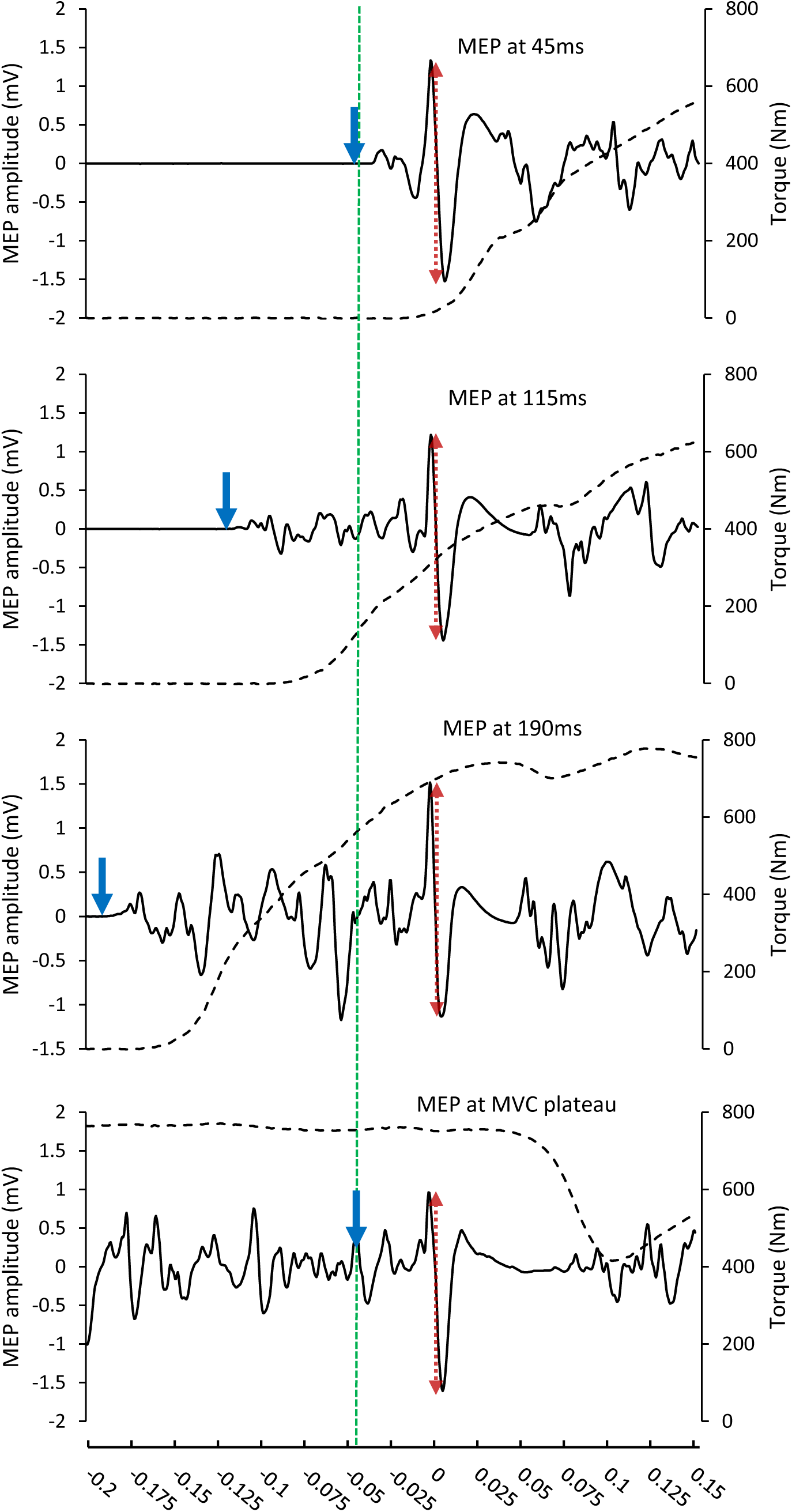
Representative traces of motor evoked potentials in the vastus medialis (solid black line) and knee extensor torque (dashed black line) for the three explosive contraction phases (top three plots) the MVC contraction phase (bottom plot). All traces are time-aligned to moment of TMS (dashed blue line). The downward-pointing blue arrows indicate EMG onset for the explosive contractions and manual TMS trigger point for the MVC contraction. The red dotted arrows represent the peak-to-peak amplitude of the MEPs used for analysis. The labels in the top three plots reflect approximate MEP times relative to EMG onset. To enhance MEP clarity, the x-axis scale switches from 50ms per tick mark pre-TMS to 30 ms per tick mark post-TMS.

**Figure 3:**
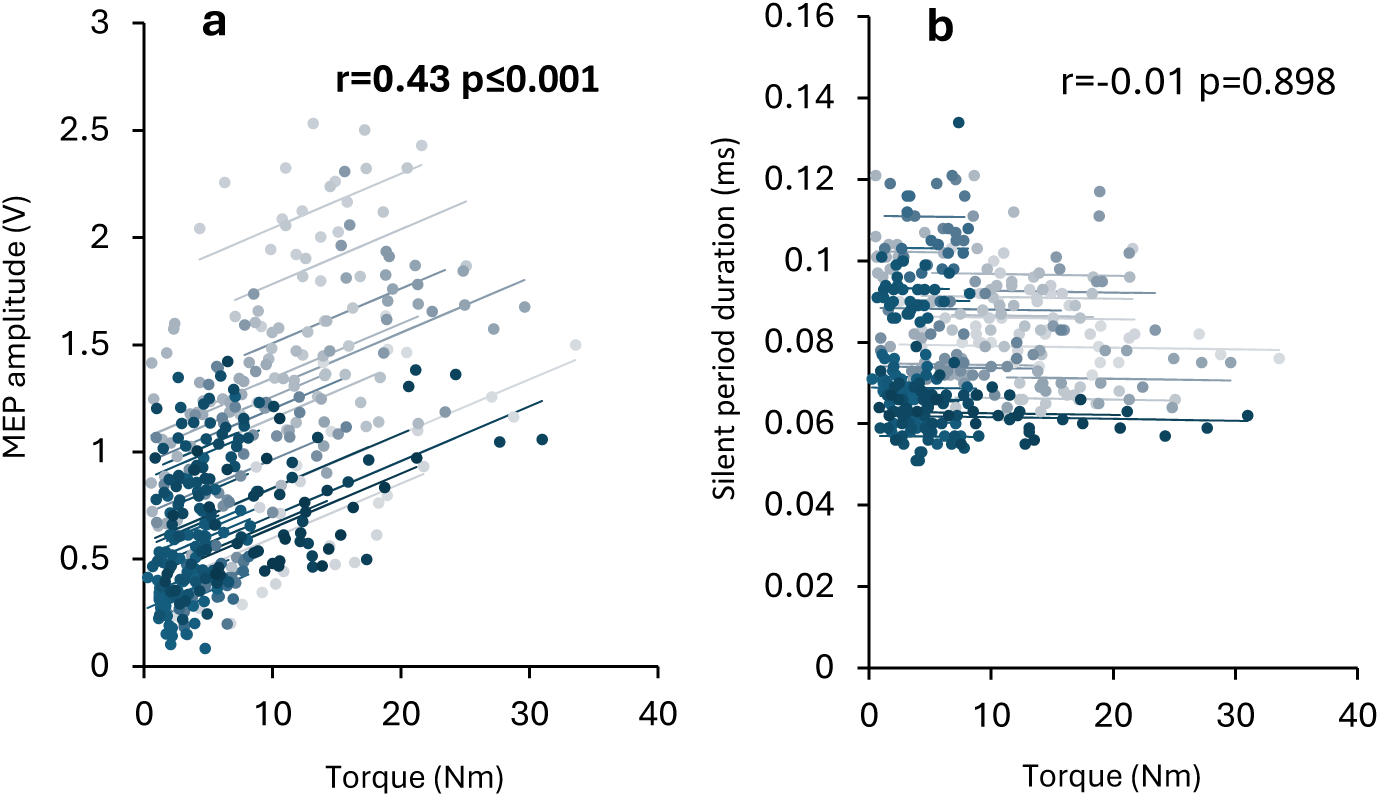
Repeated measures correlations between absolute explosive torque and TMS responses in the early phase of explosive contractions. TMS responses are absolute MEP amplitude (a), MEP relative to background EMG (MEP/EMG; b), and silent period (c). Background EMG amplitude is the RMS EMG recorded in the 20 ms preceding each MEP. Torque was recorded at 30 ms from torque onset in the same contractions as the MEP. The dots represent individual contractions. The overall dataset is split into 28 colour-coded sub-datasets (each comprising a single session from a single participant). The lines represent the RmCorr fit applied to each sub-dataset. The correlation coefficients and P-values are also provided.

### Within-person correlation analysis

Repeated measures correlations (RmCorr) (Bakdash and Marusich, 2017) were used to assess the within-person relationships between early-phase TMS responses (absolute MEP amplitude and silent period duration) and torque measured at 30 ms after torque onset within the same explosive contraction. These correlations were only considered for the early phase of explosive contractions, and not later phases, because later phase torque would be influenced by muscle activation in the earlier phases and not just in the same phase (middle or late) as the MEP measurement for that contraction (Folland, Buckthorpe and Hannah, 2014). Thus, any results of correlations between middle or late phase TMS responses and torque would potentially be misleading as we would not know the TMS responses of earlier phases within the same contraction. To prevent between-session variance influencing within-person correlation we treated data from the two sessions as independent. As a result, each repeated measures correlation contained 28 datasets (2 datasets per participant). RmCorr analyses were performed using RmCorrShiny: https://lmarusich.shinyapps.io/shiny_rmcorr/ (Marusich and Bakdash, 2021). RmCorr were interpreted as negligible 0≤r≤0.09, weak 0.1≤r≤0.3, moderate 0.4≤r≤0.69, strong 0.7≤r≤0.89, very strong 0.9≤r≤1 (Schober and Schwarte, 2018).

### Across-participants correlation analysis

Pearson’s r bivariate correlations were used to assess the relationships between torque and TMS responses (MEP normalised to Mmax and the silent period duration) in each contraction phase across the group of participants. Torque was measured at 45 (early), 115 (middle), and 190 ms (late) after torque onset in explosive contractions without TMS and averaged across the three contractions exhibiting the highest peak slopes (Tillin *et al*., 2010). The time points for torque corresponded to the time points from EMG onset at the centre of the MEP for each phase (early, middle, and late). TMS responses were averaged across the first ten available contractions for the reasons discussed above in ‘Data screening and analysis’. To account for muscle activation in earlier phases affecting torque in later phases (Folland, Buckthorpe and Hannah, 2014), early and middle phase TMS responses were averaged for correlation with middle phase torque, and early, middle and late phase TMS responses were averaged for correlation to late phase torque. Further, torques and TMS responses were averaged across sessions 1 and 2 before correlating. Pearson’s r bivariate correlations were completed using SPSS v.28. Pearson’s correlations were considered: weak r<0.3, moderate 0.3≤r<0.5, strong 0.5≤r<0.8, and very strong r≥0.8 (Hopkins, 2000).

### Analysis of contraction phase effects

For each phase of contraction (early, middle, late, MVC plateau), TMS responses (absolute MEP amplitude, MEP normalised to Mmax, MEP/EMG, and silent period duration) were averaged over the same 10 useable contractions as the bivariate correlations (see above), within each muscle and each session. The effects of contraction phases, session and muscle were investigated via a 3-way repeated measures ANOVA. For a main or interaction effect, paired comparisons were made using post-hoc t-tests with a stepwise Bonferroni correction. The median absolute deviation (MAD) process was used to detect outliers for all variables and datasets, which have been proven to be the most robust scale measure in the presence of outliers, and the most conservative value (MAD=3) was used to define outliers (Leys *et al*., 2013). Following the process described, one participant (P14) was considered an outlier for the silent period duration, and his data were removed from further analysis of the silent period duration. Statistical analysis was done using SPSS v28 (IBM Inc., Armonk, NY). The significance level was set at p≤0.05. Data are presented in the results as mean ± SD.

## Results

### Within person correlation

Repeated measures correlation based on individual data sets showed a moderately significant correlation between early phase (45ms) absolute torque and absolute MEP amplitude (Fig.3a). However, early phase absolute torque was not correlated with silent period duration (Fig.3b).

### Across participants correlation

Normalised explosive torque and normalised MEP amplitude were weakly correlated in the late phase (Fig.4c) but moderately correlated in the early (Fig.4a) and strongly in the middle phases (Fig.4b), with the latter being statistically significant. Correlations between normalised explosive torque and silent period duration were weak and non-significant for all contraction phases (Fig.4d-f).

**Figure 4:**
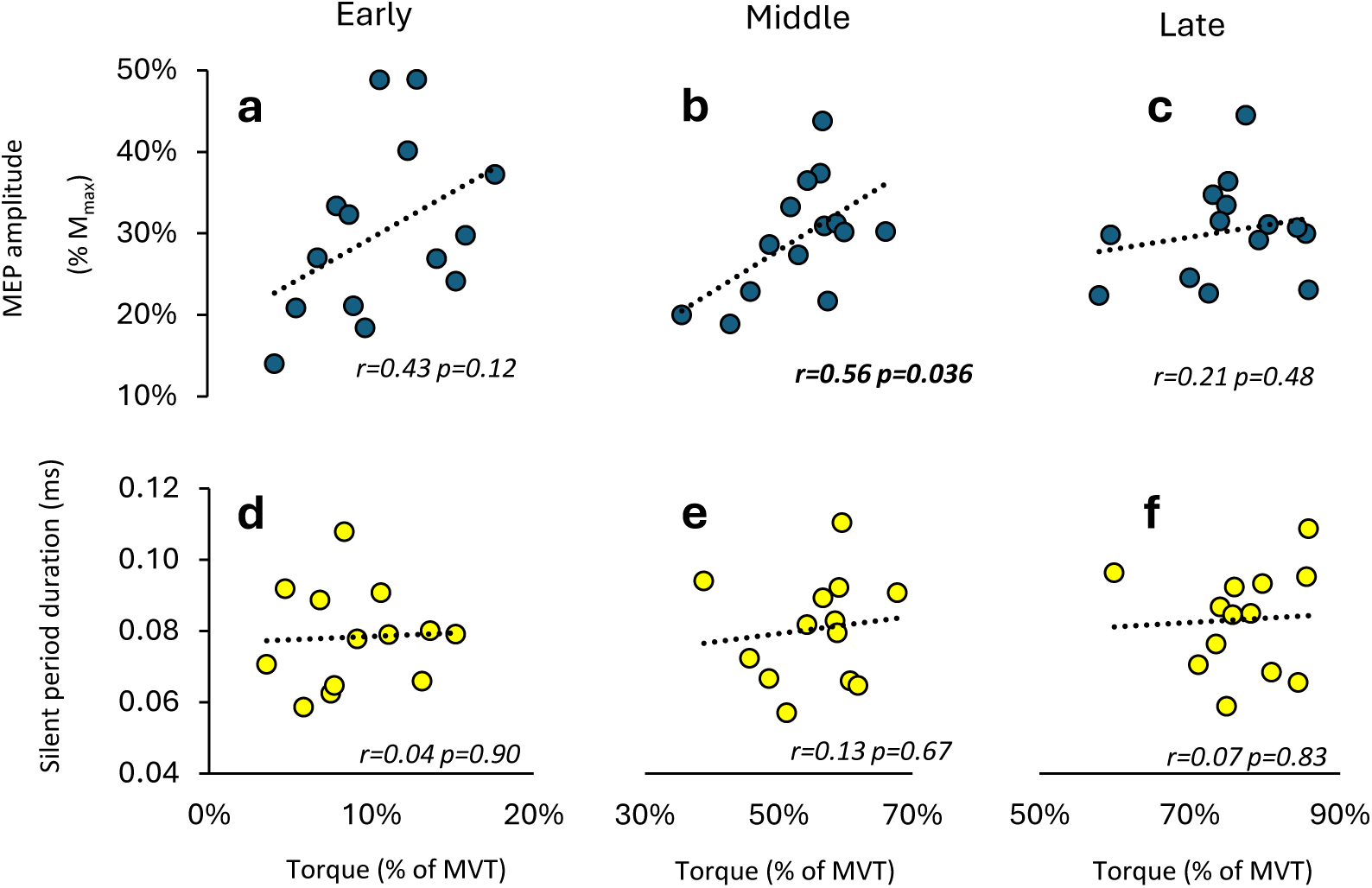
Group correlations between normalised explosive voluntary torque and TMS responses during different phases of explosive voluntary contractions. TMS responses were MEP amplitude normalised to M_max_ (a-c), MEP relative to background EMG amplitude (d-f), and silent period (j-l). Explosive torque was normalised to maximal voluntary torque (MVT). Pearson’s correlation coefficient and P-values are also presented. The phases of explosive voluntary contraction were early (45ms), middle (115ms), and late (190ms) from EMG onset (for MEP) or torque onset (for torque). Data are averaged across the three superficial quadriceps and two separate sessions for n = 14, silent period n=13. TMS responses in the middle and late phases are an average of data from that phase and the previous phase/s.

#### Effects of session, muscle, and contraction phase

No significant effects of session or any interaction effects involving session were present (Table 1). There were no observed effects of contraction phase on MEP amplitude or MEP normalised to Mmax (Fig.5c-d and Table 1). Significant effects of contraction phase were observed for MEP/EMG ratio (Fig.5e), which was higher in the early phase compared to all other phases (p≤0.001), but similar between all other phases (p≥0.9). Additionally, significant effects of contraction phase were present for the silent period duration (Fig.5f), which was shorter in the early phase compared to the late phase (-10%; p<0.001) and MVC plateau (-15%; p=0.006). Also, the silent period duration was shorter in the middle phase than in the late phase (-5%, p=0.034).

**Figure 5.**
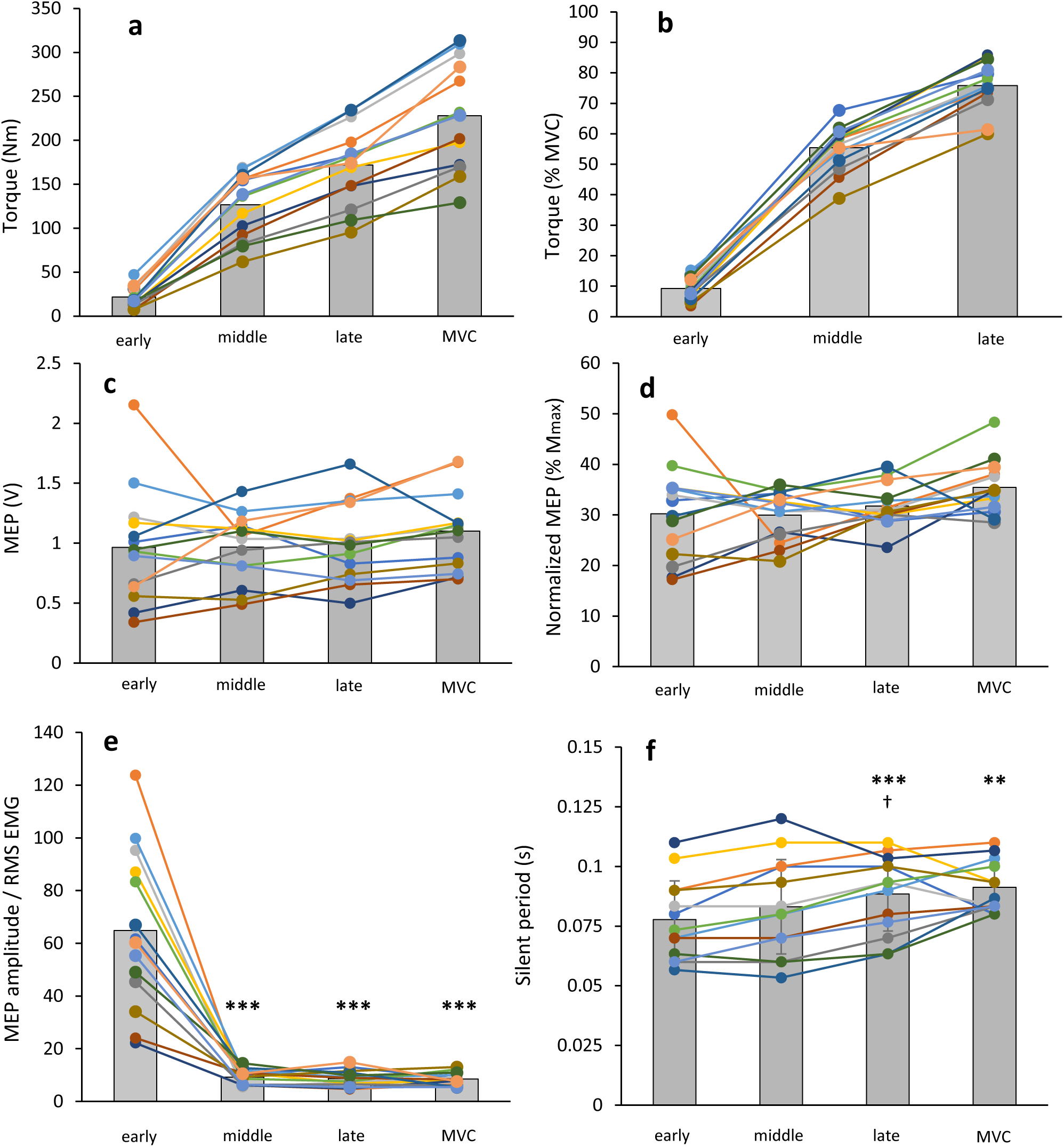
shows the effect of the contraction phase on absolute torque (a), torque normalised to MVT (b), absolute MEP amplitude (c), MEP amplitude normalised to M_max_ (d), MEP amplitude relative to background EMG (MEP/EMG) (e), and silent period duration (f). Background EMG is the RMS EMG recorded in the 20 ms from the TMS stimulation. Separate-coloured lines and dots show data sets from individual participants, while the grey bars represent group averages (n = 14, 13 for the silent period). Data were recorded at the plateau of MVCs and during the early (45 ms), middle (115 ms), and late (190 ms) phases of explosive contractions from rest. TMS response data are averaged across the three superficial quadricep muscles. Paired differences are denoted by *** (<0.001) or ** (<0.001) for different to early and † (<0.05) for different to middle.

**Table 1.** P-values of 3-way ANOVAs assessing the influence of session, contraction phase, and muscle on MEP amplitude (MEP), EMG amplitude, and silent period. In absolute terms, MEP is normalised to maximal M-wave (M_max_) and relative to background EMG amplitude (MEP/EMG). Background EMG amplitude is the RMS EMG recorded in the 20 ms preceding each MEP. These variables were measured in two separate sessions, at 3 phases of explosive contraction (early, middle, late) and the MVC plateau, and in 3 separate quadriceps muscles (RF, VL, and VM).

| Main Effects | Absolute MEP | MEP as<br>% $M_{max}$ | MEP/EMG | Silent Period |
| --- | --- | --- | --- | --- |
| Session | 0.168 | 0.853 | 0.722 | 0.88 |
| Contraction phase | 0.421 | 0.069 | **<0.001** | **<0.001** |
| Muscle | **<0.001** | **<0.001** | 0.088 | **<0.001** |
| Interactions |  |  |  |  |
| Session*Contraction phase | 0.507 | 0.521 | 0.619 | 0.875 |
| Session*Muscle | 0.588 | 0.834 | 0.201 | 0.208 |
| Contraction phase *Muscle | 0.720 | 0.413 | 0.311 | **<0.001** |
| Session* Contraction phase *Muscle | 0.488 | 0.147 | 0.268 | 0.352 |

There were effects of muscle on MEP amplitude (absolute and normalised to Mmax; Table 1), with greater absolute MEP amplitude in the VM than VL or RF (p < 0.001), but similar between VL and RF (p = 0.99). Conversely, MEP amplitude normalised to Mmax was greater in the RF than the VM (p = 0.003) and VL (p = 0.013) but similar between the VM and VL (p = 0.99). For the silent period duration, there was a main effect of muscle (Table 1) due to silent period duration being longer in the RF compared to VM (+0.004 ms, p=0.005) and VL (+0.004 ms, p=0.005), but similar between VM and VL (p=0.99).

A contraction phase*muscle interaction effect was observed only for the silent period duration (Table 1). In the early phase, the silent period duration was shorter than the late phase in all muscles (p<0.002) and shorter than the MVC phase in all muscles (p≤0.04; Fig.6). Additionally, the silent period duration was shorter in the middle phase than the late phase in the VM (p=0.02) and RF (p=0.02), and shorter in the middle phase than MVC plateau in the RF (p=0.014; Fig.6).

**Figure 6.**
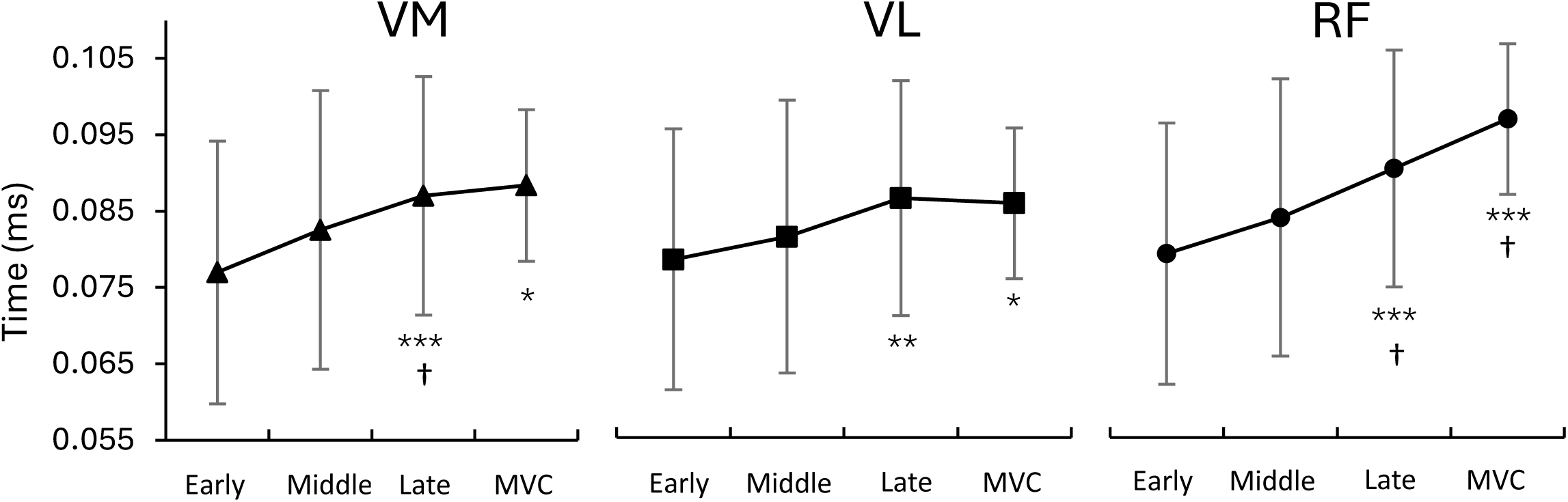
The interaction between contraction phase*muscle on silent period duration. The muscles were vastus medialis (VM), lateralis (VL), and rectus femoris (RF). Data are mean ± SD. Data were recorded at the plateau of MVCs and during early (45 ms), middle (115 ms), and late (190 ms) phases of explosive contractions from rest. Paired differences are denoted by * (p<0.05), ** (p<0.01), *** (p<0.001) for > early; and † (p<0.05) for > middle.

## Discussion

We observed positive correlations between MEP amplitude and explosive torque production in the early to middle phase of contraction, both between contractions within a person and across participants. This provides novel evidence that corticospinal excitability plays a critical role in RTD generation. In addition, we investigated MEP amplitude throughout the different phases of explosive contraction and at the MVC plateau. We found that MEP normalised to Mmax did not change in these different contraction phases, contrasting our hypothesis that it would be highest in the early phase. However, greater MEP/EMG in the early phase compared with later phases was consistent with our hypothesis, and suggests that high corticospinal excitability at the start of an explosive contraction does not immediately manifest as high EMG activity, but rather precedes it. By later phases, EMG has risen and changes proportionally with corticospinal excitability from that point. Also consistent with our hypothesis, the silent period duration increased with time and tension after contraction onset, being shortest in the early phase, followed by the middle and late phases, and finally MVC plateau. These changes in silent period duration suggest corticospinal inhibition was low in the early phases but increased with torque development over time throughout an explosive contraction.

### Association between Torque production and TMS responses

This study is the first to report a correlation between MEP amplitude and RTD across separate contractions within a person and across participants. These correlations were most observable for the early phase for within-person correlation and the middle phase across participants and suggest corticospinal excitability is an important determinant of RTD in the early and middle phases of contraction. It is well established that RTD in the early to middle phase of contraction are correlated with muscle activation (De Ruiter *et al*., 2007; Folland, Buckthorpe and Hannah, 2014; Del Vecchio *et al*., 2019), and our results imply greater corticospinal excitability is required to maximise muscle activation and thus RTD in the early and middle phases.

We can only speculate as to why the across-participant correlation between MEP amplitude and torque was significant in the middle, but not the early phase of explosive contraction, despite previous studies reporting stronger correlations between muscle activity and RTD during the early phases (Folland, Buckthorpe and Hannah, 2014). As the middle phase MEP represents an average of both the early and middle phases, thus reflecting excitability across both phases, this averaging process likely improved the reliability of the MEP data, making it more likely to observe a correlation across participants. Additionally, RTD and MEP amplitude measures are more repeatable in middle than early phases(Folland, Buckthorpe and Hannah, 2014; Castelli *et al*., 2024), which would also improve the likelihood of observing a correlation in the middle phase in our study. Finally, our sample size (n=14) was smaller than other studies correlating neurophysiological variables to RTD (n≥20, Folland, Buckthorpe and Hannah, 2014; Del Vecchio *et al*., 2019). Thus, we may have observed stronger correlations between MEP and RTD had we been able to test a greater sample size; however, this was unfortunately not possible because of disruptions caused by the COVID crises.

The silent period duration was not correlated with RTD in any contraction phase, either within- or across participants. However, when variability is low and mean values are similar across participants, the potential for significant correlations diminishes. Using the same methods, we have previously observed low variability and similar mean values in silent period durations within and between individuals (Castelli *et al*., 2024), which might reduce the likelihood of detecting correlations. Nevertheless, the silent period durations we observed during the explosive contractions were shorter (78-91 ms) than those observed by others (>100 ms) during constant load contractions and MVCs (Säisänen *et al*., 2008; Gruet *et al*., 2013). Thus, corticospinal inhibition appears negligible during an explosive contraction and is not an important determinant of RTD.

### Effects of contraction phase on corticospinal excitability and inhibition

This is the first study to examine the influence of contraction phases during explosive contraction on MEP amplitude and silent period duration. Contrary to our hypothesis, MEP amplitude (absolute or normalised to Mmax) was not highest in the early phase of EMG activity but remained constant throughout the different phases of contraction and up to the plateau of MVC. Thus, corticospinal excitability for the agonist muscles appears constant throughout an explosive contraction and is comparable to that at MVC plateau. This contrasts with the observations of Mackinnon and Rothwell (2000) for dynamic actions, where MEP amplitude was observed to peak in the early phase of agonist activity, just prior to movement onset, while decreasing during the movement. However, these dynamic rapid movements were characterised by a triphasic agonist-antagonist-agonist activation pattern, with agonist corticospinal excitability first increasing, then decreasing for the antagonist phase, and then increasing again during the second agonist phase (Mackinnon and Rothwell, 2000). The constant corticospinal excitability we observed appear unlikely to explain the dynamic modulation of muscle activation during explosive contractions, previously observed by Tillin, Pain and Folland (2018), where muscle activation increases to 120% of its level at the MVC plateau, during the early to middle phase of explosive contraction, before decreasing and plateauing at 100%. We propose these dynamic changes in muscle activation are instead driven by corticospinal inhibitory mechanisms, which we discuss further below.

Whilst MEP amplitude did not differ between contraction phases the MEP/EMG was considerably greater in the early phase compared to all other phases, but similar between middle phase, late phase and MVC plateau (Fig. 5a). Thus, corticospinal excitability increases disproportionately with muscle activation in the early phase of contraction but changes proportionally with muscle activation at later phases and at the MVC plateau. Similar effects have been observed during rapid dynamic wrist movements by Mackinnon and Rothwell (2000), who suggested the greater MEP/EMG in the early phase of EMG activity was due to increased subthreshold excitability at a cortical level, preceding spinal and then muscle activation. The authors ruled out the increases in MEP/EMG being caused by changes at a spinal level because the H-wave/EMG ratio remained constant throughout all contraction phases. We therefore speculate that cortical excitability increases ahead of muscle activation at the start of an explosive contraction, and it takes until the middle phase for muscle activation to ‘catch-up’ and change proportionately with cortical excitability thereafter. The increase in subthreshold cortical excitability at the start of an explosive contraction likely commences up to ∼70 ms prior to EMG onset (Baudry and Duchateau, 2021).

Our results provide novel evidence that the silent period duration generally increased throughout an explosive contraction and up to MVC plateau, with it being lower in the early phase than the late phase and MVC plateau, and lower in the middle phase than the late phase (Fig.5f). Shorter silent period durations in the early phase supports the notion that inhibition at a spinal and supraspinal level is low to maximise muscle activation and RTD during an explosive contraction. However, corticospinal inhibition appears to increase with increased torque development during the explosive contraction and is highest at MVC plateau. The mechanisms underpinning this effect are unclear, but the silent period durations may provide some clues (Škarabot *et al*., 2019). The short silent period durations we observed during the different phases of explosive contraction (78-90 ms) indicate inhibitions at a spinal level (rather than supraspinal) are driving variability in silent period. Contributing factors to these spinal inhibitions are likely to include some motor units being in a hyperpolarised state at the time of TMS, Renshaw cell inhibition, reciprocal inhibition, and di-synaptic inhibition (Triggs *et al*., 1993; Ziemann *et al*., 1993; Enoka and Fuglevand, 2001), all caused by activation of the motor units and/or torque development in the early phase of contraction. In contrast, the silent period durations towards the later phase of explosive contraction (86 ms) and at MVC plateau (92 ms) indicate cortical mechanisms, likely involving intracortical inhibition due to GABA B receptors activity (in addition to spinal inhibitions) are contributing to silent period length (Werhahn *et al*., 1999; McDonnell, Orekhov and Ziemann, 2006). The lower corticospinal inhibitions in the early phase of contraction followed by increased inhibition at later phases and at MVC plateau, potentially explains previous observations of muscle activation being greater in early to middle phases of explosive contraction compared to late phase and MVC plateau (Tillin, Pain and Folland, 2018).

### Limitations

It is possible MEP amplitudes measured in our study did not change proportionally with corticospinal excitability, which could mask possible differences between the phases. Evidence for this is based on studies showing that MEP amplitudes increase with increases in constant force output up to 55-75% MVC and then plateau or slightly decrease beyond this intensity (Todd, Taylor and Gandevia, 2003; Goodall, Romer and Ross, 2009; Škarabot *et al*., 2018). This is likely attributable to the high motoneuron discharge rates observed during maximal effort (explosive and MVC) contractions. Under these conditions, there’s an increased likelihood that many motor units will be in their refractory phase, reducing the probability of them responding to TMS. Nevertheless, we did observe sufficient variability in MEP amplitudes both within and between people, to enable correlations between MEP amplitude and RTD. Therefore, it is conceivable our methods were sensitive enough to measure differences in MEP amplitude between phases, had they existed.

We cannot empirically establish the specific site (e.g., cortical, intracortical, or spinal) of excitation and inhibition in the different contraction phases with TMS alone. We recommend that future studies use additional approaches (e.g., spinal stimulation procedures or methods that capture subthreshold rise in corticospinal excitability as for Mackinnon and Rothwell, 2000) to explore our observations further.

## Conclusion

MEP amplitude was correlated with RTD over the early phase (within people) and middle phase (across participants) of explosive contraction, suggesting corticospinal excitability is an important determinant of RTD. Although MEP amplitude was constant throughout the phases of an explosive contraction, and at the plateau of MVC, MEP/EMG was higher in the early phase, suggesting the high corticospinal excitability in the early phase is subthreshold for full muscle activation, but this likely facilitates high muscle activation and, in-turn, torque output at later phases. The silent period duration was lowest in the early phase and increased throughout the different phases, suggesting that corticospinal inhibition increases with increased torque during an explosive contraction and appears highest at the MVC plateau.

The authors have no relevant financial or non-financial interests to disclose.

## Acknowledgements

The authors would like to sincerely thank Boston Freestone for its vital contributions to data acquisition.

## Abbreviations

AMT: Active motor threshold
RMT: Resting motor threshold
GABA: Gamma-aminobutyric acid
EMG: Electromyography
MEP: Motor evoked potential
MVC: Maximal voluntary contraction
MU: Motor unit
RF: Rectus femoris
RmCorr: Repeated measures correlation
RMS: Root mean squared
RTD: rate of torque development
SD: Standard deviation
SP: Silent period
TMS: Transcranial magnetic stimulation
VM: Vastus medialis
VL: Vastus lateralis

## Notes

### Competing Interest Statement

The authors have declared no competing interest.

